# Body-like coordination emerges in paired termite locomotion

**DOI:** 10.64898/2026.05.22.727143

**Authors:** Nobuaki Mizumoto, Sayomi Kamimoto

**Affiliations:** Department of Entomology & Plant Pathology, Auburn University, Auburn, AL, 36849, USA; Department of Mathematics, Howard University, Washington, D.C., 20059, USA

**Keywords:** active sampling, leadership, locomotion, superorganisms

## Abstract

Coordinated group movements are often described as moving “like a single organism,” yet this analogy is typically a metaphor. Individuals, interacting with each other, lack the functional integration of the body parts of a single organism. It remains unresolved whether movement coordination can truly reproduce body-like organization. Here we show that during tightly coordinated movement, pairs of termites use the same exploration–stabilization division of labor observed within a single moving body. Using posture tracking, we demonstrate that leader– follower asymmetries in tandem running mirror anterior–posterior asymmetries in individual locomotion, with exploratory motion concentrated at the front and smoother, shorter trajectories at the rear. Our mathematical model reveals that such body-like coordination emerges from hierarchical interactions between leaders and followers, described by the temporal convolution of past locational cues rather than instantaneous responses. These results identify a control principle of hierarchical structured movement coordination, providing a novel way to design collective behavior.

## Introduction

A central goal of biology is to identify principles that operate across scales. In biological systems, higher-level organization emerges through the aggregation and functional differentiation of components (1), from cells to organisms, and from individuals to collectives (2–4). As in bird flocks, fish schools, and insect swarms, coordinated movement in animal groups is often described as resembling that of a single organism (5, 6). Collective behavior emerges from coordination among individuals (7, 8), whereas animal locomotion arises via coordination among body parts within an individual (9–11). In both cases, multiple components operate in a distributed manner while producing integrated movement as a whole (9). Despite these similarities, there is no general framework that connects coordination within individuals to coordination among individuals, leaving the resemblance largely metaphorical.

Within individuals, movement coordination among body parts is inherently asymmetric along the body axis. Environmental exploration by blind animals is driven by active sampling (12, 13); individuals swing the sensory units in anterior body regions (e.g., antennae and heads), while posterior regions maintain cohesion (11, 12, 14–16). Similarly, coordinated animal groups often exhibit leader-follower structures; leaders sample the environment, and followers constrain their motion to preserve group integrity (17, 18). Although the leader-follower relationship may emerge from small differences in individual properties (19), leadership is structurally inevitable in contact-based communication that typically exhibits single-file collective motion (20). Because sensory organs are typically located anteriorly, each individual follows the one ahead by maintaining physical contact (21–26). In such systems, interactions are directional from front to back, analogous to the coordination of body parts for locomotion. We hypothesize that both within- and across-individual coordination arise from the same principle of role separation between exploratory and stabilizing components along a chain.

Mating termites offer an ideal system for linking within- and across-individual movement coordination. During the mating season, individual termites search for a mating partner, and paired females and males form a tandem running while seeking sites for colony foundation (27). Termites rely on physical contact for exploration, in which searchers actively swing their antennae to find partners (28, 29). Similarly, tandem running in termites is maintained by contact-based communication and directional interactions (28–31). The female leads the tandem while actively sampling the environment to find a nesting site, and the male keeps touching the female’s abdomen with its antennae and mouthparts (29, 32). In physical contact, information flow is from the leader to the follower; the leader predominantly determines the course and timing of movement (30, 33). In both situations, there is a functional division of labor between anterior parts (head or leader) and posterior parts (body or follower); the anterior samples the environment, while the posterior maintains integration.

To bridge whole-body locomotion and leader-driven collective motion, here we show that both systems can be understood as assemblies of agents (body parts or individuals) with functionally distinct roles connected through directional interactions. Using posture tracking, we show that both single and paired termites exhibit similar movement trajectory patterns; anterior components (heads or leaders) generate exploratory trajectories and posterior components (bodies or followers) generate smoother, stabilizing motion. We further develop a mathematical framework based on functional differentiation, in which exploratory agents independently explore the environment, while stabilizing agents maintain cohesion through order-dependent interactions. This framework provides a unified description of movement coordination across biological scales.

## Results and Discussion

### Comparison of within- and across-individual movement coordination

We tracked the movement trajectories of the body centers of leaders and followers in tandem running termites, while head tips and abdominal tips of single searching termites in two species, *Coptotermes formosanus* and *Reticulitermes speratus* (Fig. 1A). The movement trajectories of anterior (leader or head) and posterior components (follower or abdomen) of single and paired termites were nearly indistinguishable from each other (Fig. 1B, Video S1). Single termites actively swayed their heads and antennae as they explored the environment, while their bodies remained stable, resulting in distinct trajectories between the heads and bodies. The posterior body trajectories were smoother and shorter than those of the anterior heads (one-sample t-test to test if the distance ratio is smaller than 1, *C. formosanus*: *t*_19_ = -22.4, *P* < 0.001, *R. speratus*: *t*_39_ = - 10.3, *P* < 0.001; Fig. 1C). The same was observed in tandem running pairs. The leader termites actively explore the environment, while the follower termites maintain a stable position behind the leader, resulting in smoother, shorter paths for the followers (one-sample t-test to test if the distance ratio is smaller than 1, *C. formosanus*: *t*_36_ = -7.51, *P* < 0.001, *R. speratus*: *t*_39_ = -4.20, *P* < 0.001; Fig. 1C). We did not find species differences (Linear mixed model, LMM, 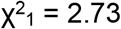, *P* = 0.10), but single termites showed smaller distance ratio than tandem running (Linear mixed model, LMM, 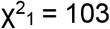, *P* < 0.001; Fig. 1C). The consistent patterns were observed in path straightness and turning angles, where anterior components show less staright paths and larger turning angles compared with posterior components (Figure S1-2). This reflects the role differentiation between the anterior and posterior components within and across individual coordinations; in a paired tandem, the leader explores the environment, while the follower maintains integration. Similarly, within a termite body, the head actively samples the environment, while the posterior body never explores independently.

**Figure 1.**
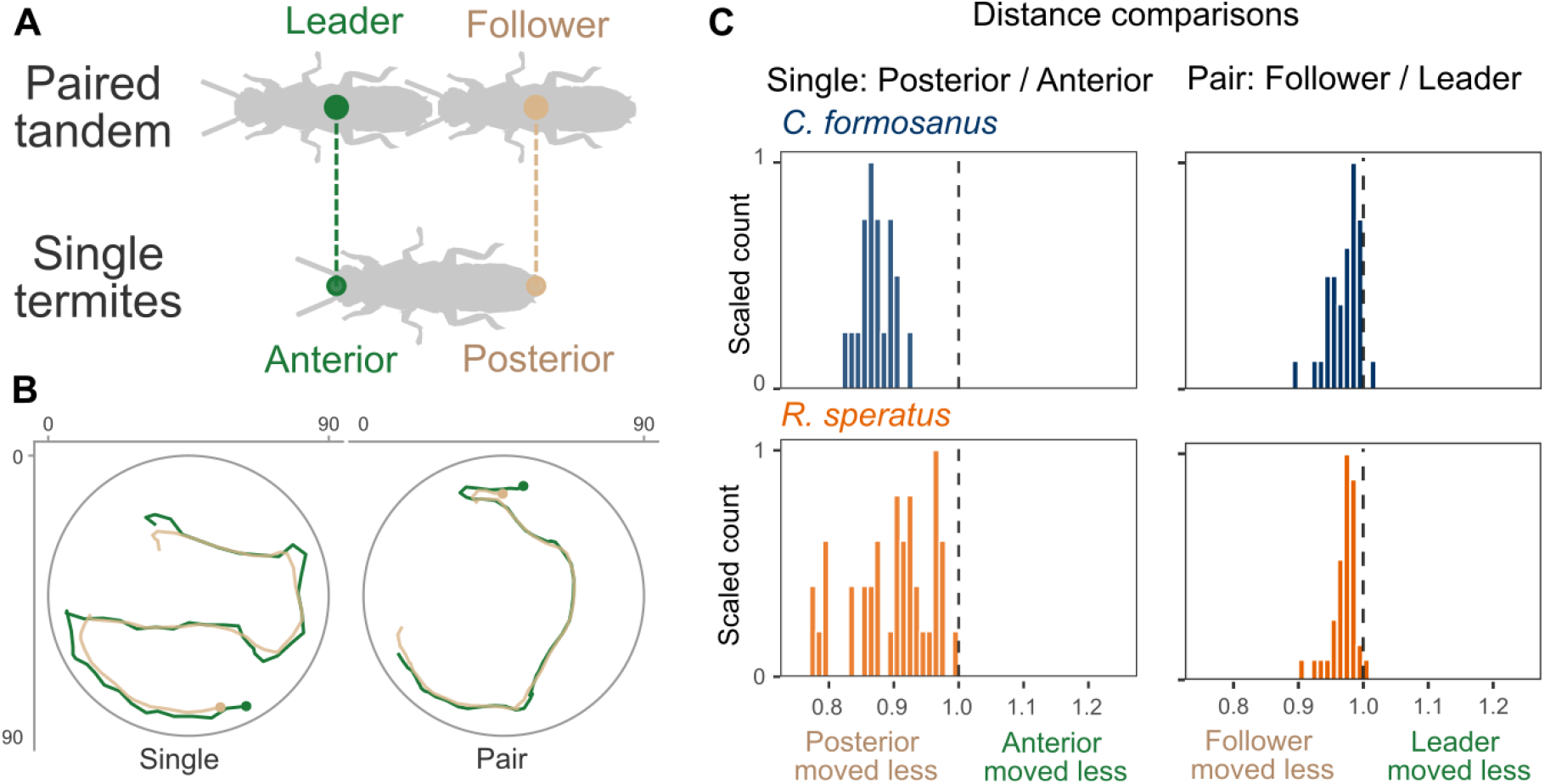
Similarity of paired movement coordination and whole-body coordination in searching termites. (A) Both paired and single termites are separated into anterior and posterior parts. (B) Anterior and posterior trajectories of paired and single termites. (C) Comparison of the leader and follower (or head and abdomen) trajectory length.

Our trajectory analysis revealed that ordered interactions and active waiting by posterior components underlie role differentiation in both paired and single-termite coordination. Step length distributions were clearly distinct between anterior and posterior components: posterior components exhibited shorter steps and more frequent pausing than leaders (Fig. 2A), indicating that shorter and smoother trajectories of posterior components also reflect pausing behaviors (Fig. 1). Accordingly, pausing behavior was asymmetric. Although movement states were usually synchronized, asymmetric states were strongly biased: anterior movement with posterior pausing occurred significantly more often than the reverse (Permutation test, *P* < 0.001 in both single and paired termites; Fig. 2B, C). During these posterior-specific pausing periods, the anterior-posterior directional changes increased, consistent with active exploration. The change in orientation was largest when anterior components moved while posterior components paused (solo: LMM, 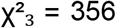, *P* < 0.001; tandem: 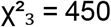, *P* < 0.001; Fig. 2D), indicating that posterior pausing coincides with anterior exploratory reorientation. This pattern is consistent with individual active sensing behaviors reported in other systems, such as *Drosophila* larvae and rodents (12, 14, 15). Our analysis revealed that the same can happen even in individual movement coordination.

**Figure 2.**
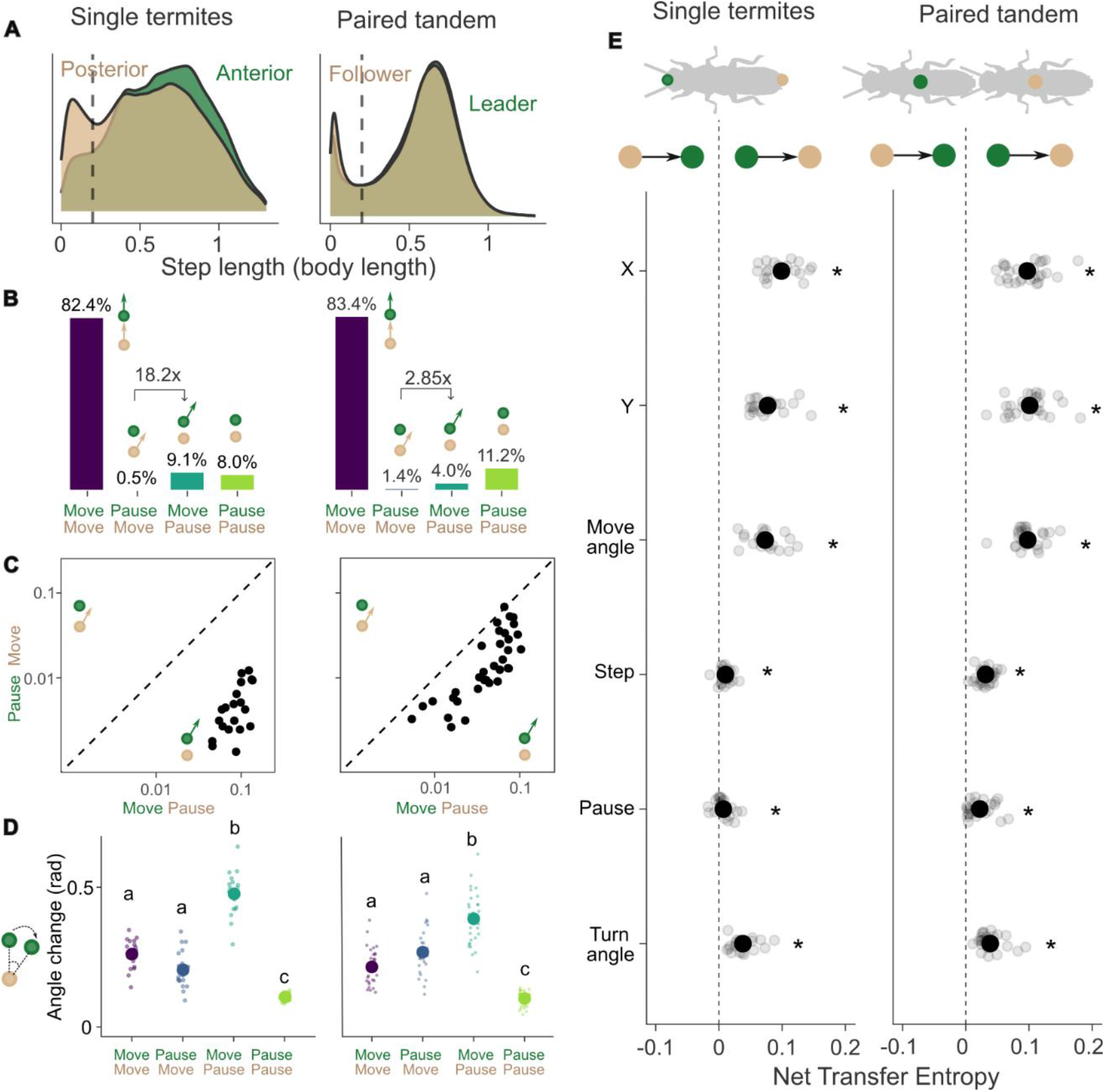
Movement coordination in paired and single termites shares the same division of labor and directional control. Data are from *C. formosanus* (*R. speratus* in Fig. S3). (A) Step length distributions show shorter and more intermittent movement in posterior components compared to anterior components, consistent with stabilizing versus exploratory roles (see Fig. S4 for breakdown). Dashed lines indicate the pausing threshold (0.2 body length). (B) Frequencies of movement states show strong asymmetry: posterior pausing while anterior moves occur more frequently than the reverse. (C) This asymmetry is consistent across individuals and pairs. (D) Turning depends on movement state and is greatest when the anterior moves while the posterior pauses, indicating exploration during posterior waiting. Different letters indicate significant differences (LMM; Tukey HSD, *P* < 0.001). (E) Net transfer entropy (TE_ante→post_ − TE_post→ante_) shows consistent directional information flow from anterior to posterior components across kinematic and behavioral variables. * indicates the statistical significance that net transfer entropy is larger than 0 (one-sample t-test).

Transfer entropy analysis further revealed consistent directional asymmetry in interactions. Information flow was predominantly from anterior to posterior across both behavioral (pausing, turning) and kinematic (step length, position) variables (Fig. 2E). Across both individual and paired systems, posterior components were systematically dependent on anterior components. Together, these results show that coordination arises from directional interactions between functionally distinct units, from anterior to posterior components. This shared structure indicates that both within- and across-individual coordination follow the same principle of role differentiation along a connected chain.

### Coordination as a directed information-processing system

Our empirical results reveal a common structure underlying both within- and across-individual coordination: interactions are directional and organized along an anterior-posterior axis, with functional role differentiation. However, most existing models of movement coordination in animal collective behaviors are based on symmetric and reciprocal interactions among anonymized agents (34–36). In such systems, all agents follow the same behavioral rules that describe their responses to estimated states of other neighbors (e.g., positions and heading directions) (8). In animals, such communication is typically mediated by vision, where spatial information is converted into effective social forces that govern movement responses (17, 37–39), successfully explaining many visually based collective phenomena (40–42). Symmetry is therefore a natural default assumption, as it enables anonymity among agents, scalability to large group sizes, and analytical tractability. Therefore, the models of collective movement coordination do not naturally account for coordination within an individual.

To bridge coordination across biological scales, a model must explicitly incorporate role differentiation and directional interactions. Such systems should achieve follower-based coordination, in which exploratory roles generate movement independently from others, while stabilizing roles maintain cohesion through order-dependent interactions driven by upstream inputs. Based on this principle, we developed a mathematical framework in which agents are arranged in a directed chain of interactions. Exploratory agents generate movement paths, whereas stabilizing agents respond to upstream cues to constrain and integrate motion. This system preserves the division of labor between exploratory and integrative components and applies irrespective of whether agents correspond to body parts or individuals.

The coordination is implemented as an order-dependent, sequential process in which interactions occur through contact points rather than through centers of mass. In contact-based communication, body shape cannot be ignored. There is a difference between a leader’s position (i.e., the body center) and the actual point where the contact happens (i.e., the leader’s posterior tip of the body) (Fig. 3A). We represent this interaction point as a leader’s proxy to distinguish it from the leader’s position. The same happens in within-body movement coordination. Insect bodies consist of segments, e.g., the head and abdomen are joined at the thorax (43), where the joint functions as a proxy for the interaction between two different body parts (Fig. 3A). The proxy is defined through a recursive temporal weighting of the exploratory component, and the integrative components update their movements moving towards the contact proxy.

**Figure 3.**
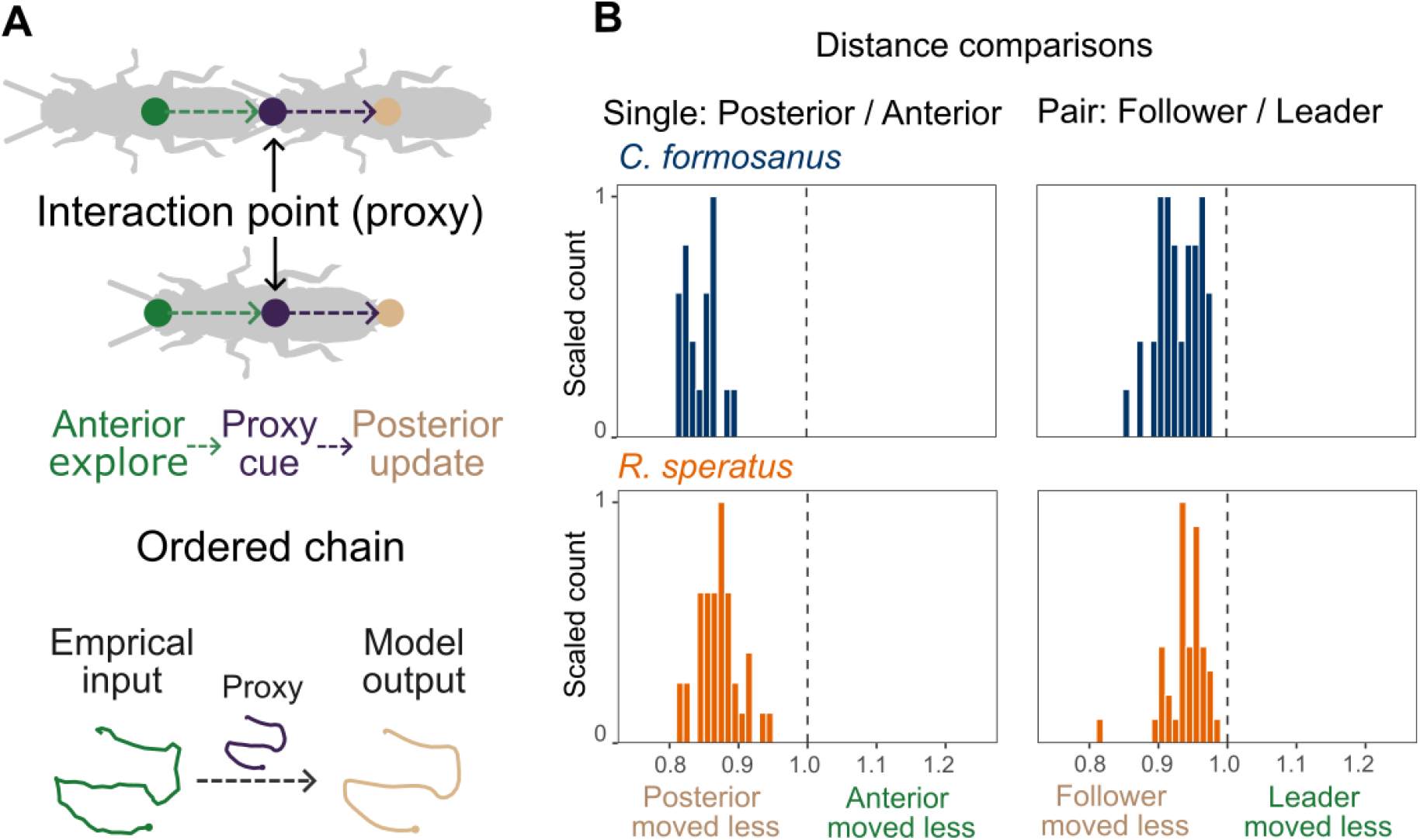
Directional model of movement coordination reproducing the observed patterns. (A) The model schematic. The anterior and posterior parts interact through a proxy rather than at their actual positions, creating an ordered chain of interactions. Empirical anterior trajectories are used to generate posterior trajectories. (B) Distributions of model-generated trajectory length compared with input empirical anterior trajectory lengths (λ = 0.38).

Despite its minimal structure, the model reproduced the key statistical features observed in both single and paired termites. Because leaders act as trajectory generators, we directly used the empirical trajectories of termite leaders or head positions as model inputs (Fig. 3A). Using the same parameter (λ = 0.38), the model generated shorter, smoother trajectories corresponding to the posterior components in both single and paired termite movement coordination (one-sample t-test, *P* < 0.001 for all indices, Fig. 3B, S6-7). Because the model does not include any noise, the model-generated trajectories were generally shorter than the empirical posterior trajectories (paired t-test, *P* < 0.001, comparing Fig. 1C and Fig. 3B) and also exhibited higher straightness and lower turning angles (paired t-test, *P* < 0.001). Our model also reproduced behaviors resembling waiting in the posterior components. The step-length distribution showed that the model-generated posterior step lengths were generally smaller than the empirical anterior trajectories, but did not exhibit clear inflated pausing patterns (Fig. S8). On the other hand, anterior-posterior directional changes occurred when posterior components moved shorter distances (Fig. S9).

### Temporal integration of the exploration path governs coordination

While the model reproduced empirical patterns using a common parameter, it remains unclear how this parameter shapes coordination dynamics. Our model is governed by a single parameter (λ) that determines how posterior components integrate anterior exploration movements over time (Fig. 4A). We defined the proxy position as **r**_**P**_(*t*) = (1 − λ)**r**_**P**_(*t* − 1) + λ**r**_**L**_(*t*), where λ controls how much past information is integrated. This parameter, therefore, determines the integration timescale: small values correspond to long integration (strong memory), whereas large values correspond to short integration (rapid tracking).

**Figure 4.**
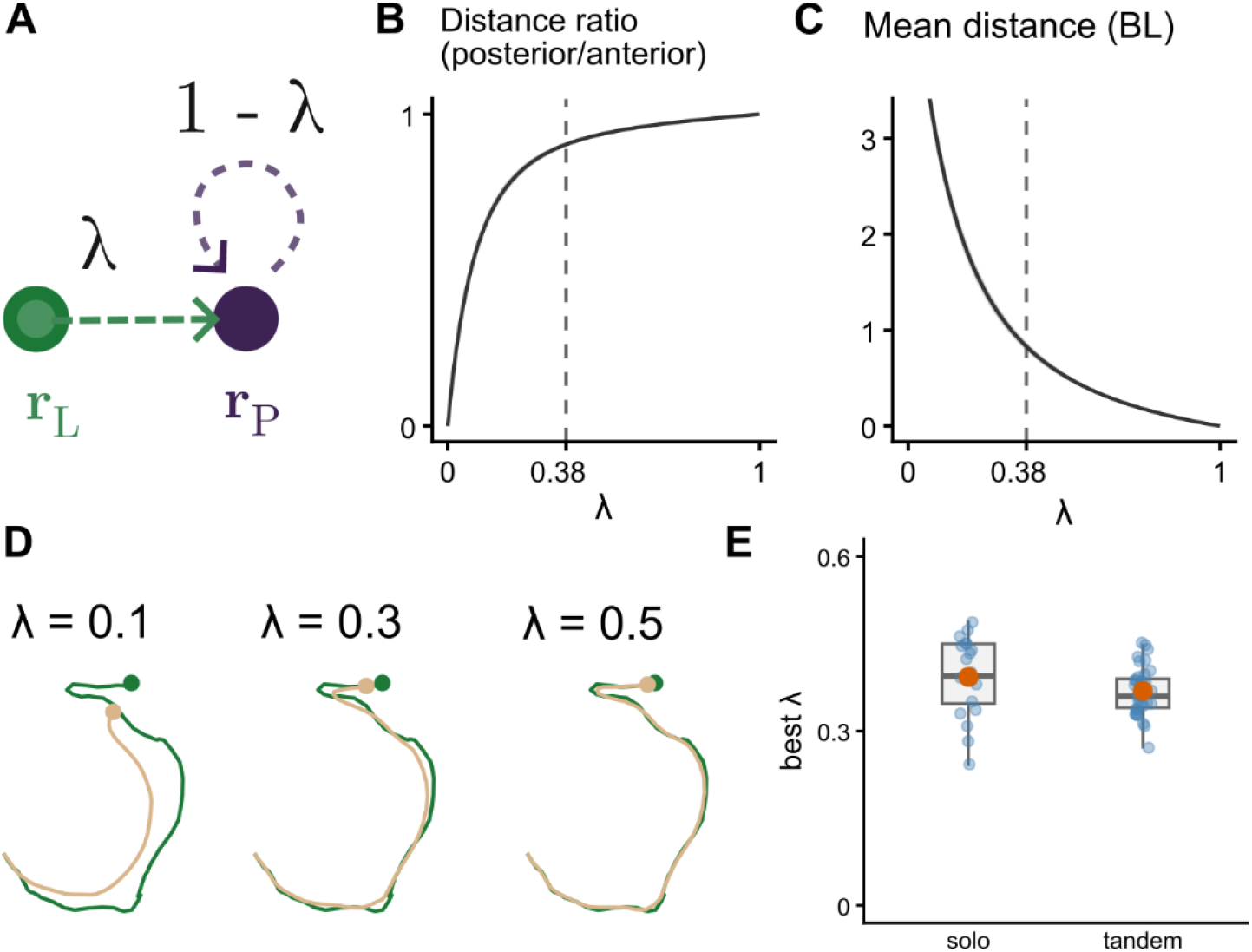
The effect of time integration parameter, λ, on movement coordination.(A)Schematic of the effect of λ on time integration, where a smaller λ leads to longer time integration. (B) The effect of λ on distance ratio. The global mean, λ = 0.38, which is used for Fig. 3, is indicated. (C) The effect of λ on the distance between posterior and anterior components, in body length units. (D) Example trajectories for different values of λ. (E) Comparison of the best fitted λ for each trajectory of single termites and paired termites in *C. formosanus*.

Change in the parameter λ revealed distinct coordination regimes. At low values, posterior components exhibit strong smoothing, leading to reduced movement, increased delays, and weaker coupling with anterior trajectories (Fig. 4B-D). At high values, on the other hand, posterior components rapidly track anterior motion, leading to tight coupling without clear functional differentiation between components (Fig. 4B-D). We found that the system produces stable coordination, similar to both within- and across-individual coordination in termites (λ ∼ 0.3; Fig. 4B-D).

The model fitting for each trajectory revealed that the estimated parameter values were similar across species and interaction scales. There was no significant difference between within-individual and across-individual coordination (t-test, *t*_26.6_ = 1.47, *P* = 0.15; Fig. 4E, S10), indicating that a common integration timescale governs coordination dynamics in both scaled systems. Note that the λ value was generally estimated to be larger in within-individual coordination, suggesting that mechanical coupling is stronger due to physical and neurological connections within the individual body (Table S2). Together, these results show that variation in a single integration parameter is sufficient to reproduce distinct coordination patterns and provide a unified explanation for movement coordination across biological scales.

### Conclusion: Unified framework connecting body coordination and group coordination

Despite extensive study of both locomotor control within organisms and collective coordination among organisms, these problems have been treated as fundamentally distinct. In contrast, our results show that these two coordinations can be described by the same mechanism of exploration-integration division of labor. In both head–body coordination and leader–follower coordination, anterior components primarily generate exploratory paths by environmental sensing, while posterior components primarily maintain geometric coherence by smoothing trajectory fluctuations. Functional division of labor among components is key; it is a natural consequence of individual locomotion in many elongated animals, where directional changes are typically initiated at the anterior part with sensory inputs and propagate to the posterior part through mechanical and neural links (44–46). Contact-based group movement, such as tandem running, also naturally reproduces this structure as individuals maintain strict spatial positioning connected by anterior-posterior physical contacts (29, 47). In non-contact communication, the relative positions of leaders and followers are more flexible, though leaders typically position themselves near the front (48, 49). Given that some groups show fixed leadership roles (50, 51), it is an open question whether the inter-individual movement coordination can represent within-individual coordination in such systems.

Importantly, contact-based communication does not automatically create a body-like movement coordination. Tandem running recruitment in *Temnothorax* ants is the most extensively studied example of contact-based movement coordination (52). In this system, tandem running primarily functions as a form of route learning rather than movement coordination, in which leaders guide naïve followers to a resource (30, 53, 54). Consistent with this function, our additional analysis on *Temnothorax rugatulus* showed a reversed movement pattern relative to body-like coordination: followers move greater distances than leaders (Fig. S11). Thus, posterior followers exhibited explorative movement associated with route learning, while anterior leaders maintained a stable path. This contrasts with other ant systems, in which tandem running primarily serves movement coordination, such as *Diacamma indicum* (55), where followers consistently exhibited shorter and smoother paths than leaders (Fig. S11). Together, these comparisons suggest that the division of labor between exploration and integration depends on the functional context of the interaction, and is not an automatic consequence of contact-based coupling.

Beyond a pair, we offer an alternative approach for modeling collective movements by providing functional asymmetry in movement coordination. Standard movement coordination models, especially in animal collective behavior, seek the mechanisms that create directional persistence within a group through inter-individual alignment or consensus (8, 56). Such systems often assume symmetric, exchangeable interactions (34–36), eliminating spatial role asymmetry. These models successfully explain the symmetric collective movement modes such as swarming, milling, and schooling (or flocking) (36, 57–59), yet structurally exclude the class of behaviors that require directional information flow and non-exchangeable roles. Single-file coordination, as in within-individual coordination or tandem running in this study, or widely observed in the animal kingdom, from extinct trilobites to extant mammals (21–26), belongs to this excluded class. Also, single-file motion can happen in any group species constrained to move in a narrow path (20). Our results demonstrate that spatio-temporal integration of upstream information by downstream components is key to directional coordination, captured by a temporal integration process of past positions, rather than instantaneous alignment rules. By expanding the idea of movement coordination, our study can also be applied in human systems that rely on directional information flow, such as in truck platoons and automated convoys (60, 61).

## Methods

### Behavioral data

Trajectory data were compiled from previously published datasets of tandem-running and solo termites (28, 29, 33). For tandem pairs, body-center trajectories were used to represent leader-follower movements, while for solo individuals, head and abdominal tip coordinates were used to characterize anterior-posterior movement. Data were obtained for *Coptotermes formosanus* and *Reticulitermes speratus* across multiple studies conducted in Japan and the USA between 2017 and 2022, where original trajectories were extracted using either UMATracker (62) or SLEAP (63) (see Table S1 for full information on all videos). In total, we obtained 37 tandem and 20 solo trajectories for *C. formosanus* and 40 tandem and 40 solo trajectories of *R. speratus*, where each trajectory corresponds to a unique individual or pair. Body length was measured for each individual in the previous studies, and all spatial coordinates were rescaled by body length (pair mean for tandem data). All datasets were temporally standardized by downsampling to every 0.2 seconds (the original recordings were 30–60 frames per second), and trajectories were standardized into a common format for analysis. All analyses were performed using R ver 4.5.2 (64).

### Trajectory analysis

For paired trajectories, termites were classified as tandem running when the distance between body centers was less than two body lengths (65). Analyses of paired coordination were restricted to these periods. For solo individuals, the full trajectories were used, as head and abdominal tip distances are inherently bounded within this range.

Distance ratio: We computed the total traveled distance covered by anterior components (head or leader) and posterior components (abdomen or follower). For each trajectory, we calculated the distance ratio as: posterior distance / anterior distance. For each species and dataset type (solo or tandem), we tested whether the mean distance ratio was less than 1 using one-sample t-tests. Normality was assessed using Shapiro–Wilk tests. The assumption was not violated for *C. formosanus* (P > 0.13), but was violated for *R. speratus* (P < 0.05). We retained parametric tests for consistency, noting that the results were qualitatively consistent with those from the nonparametric Wilcoxon signed-rank test. The same protocol was used for the subsequent one-sample t-tests as well. We also compared distance ratios across species and dataset types by using a linear mixed-effects model (LMM) with distance ratio as the response variable, species and dataset type as fixed effects, and individual or pair identity as a random effect. We used the lmer() function from the R package ‘lme4’ (66) and tested statistical significance with a type II ANOVA using the Anova() function in the ‘car’ package.

Path straightness: We tested whether the posterior paths were smoother in two approaches. First, we measured the straightness of the path, defined as the ratio of net to gross displacement (67). Along each trajectory, net displacement was defined as the Euclidean distance between positions at frames *t* − 3 and *t* + 3, and gross displacement as the cumulative path length across the same interval (using alternative interval lengths yielded qualitatively consistent results). Straightness was averaged across frames to obtain a trajectory-level measure. Then, straightness was compared between anterior and posterior trajectories. Second, we measured the turning angles of each trajectory as the magnitudes of changes in direction of motion between successive frames. The turning angles were obtained only when the components were not pausing (see below). We obtained the mean absolute turning angles (in radians) as a measure of path sinuosity and compared the anterior and posterior trajectories using t-tests.

Pausing dynamics: Step lengths between successive frames (0.2 s) showed a bimodal distribution (Fig. 2A, S3A, S4-S5), corresponding to pauses and movements. We defined a threshold at 0.2 body lengths/sec, corresponding to the local minimum between the two modes of distribution. Displacements below this threshold were classified as pauses. We examined other threshold values, but they did not qualitatively affect our conclusions. Using anterior and posterior move/pause states, we defined four coordination states: move–move, move–pause, pause–move, and pause–pause. We quantified coordination asymmetry by comparing the frequencies of move– pause and pause–move states. The asymmetry index was defined as: log((move–pause) / (pause–move)). To test whether the observed asymmetry differed from chance, we performed permutation tests by randomly shuffling the temporal order of posterior pausing states (1,000 iterations). P-values were calculated as the proportion of permuted datasets with asymmetry indices greater than or equal to the observed value.

Angular dynamics across pausing states: We computed the change in relative angle (radians) between the posterior and anterior components at each time step (0.2 s) and calculated the mean absolute change in relative angle for each coordination state. Differences among coordination states were tested using LMMs, with coordination state as a fixed effect and individual or pair identity as a random effect. Statistical significance was assessed using type II ANOVA with the Anova() function in the ‘car’ package, followed by Tukey post hoc tests with the glht() function in the ‘multcomp’ package (68). Trajectories in which asymmetric states (move–pause or pause– move) comprised less than 1% of observations were excluded from this analysis.

### Information flow

We quantified directional information flow between anterior and posterior components using transfer entropy, implemented with the transfer_entropy() function in the ‘RTransferEntropy’ package (69). Transfer entropy measures the reduction in uncertainty of one time series given the past state of another (70). Thus, transfer entropy is asymmetric between two time series; from anterior to posterior and from posterior to anterior. Reliable estimation of transfer entropy requires sufficiently long time series. Based on the previous power analysis (30), we used continuous sub-trajectories of tandem running events, lasting more than 10 minutes (>3,000 steps) (Fig. S12). We obtained 29 tandem and 20 solo trajectories for *C. formosanus* and 30 tandem and 40 solo trajectories of *R. speratus*. We computed transfer entropy in both directions (anterior → posterior and posterior → anterior) for the following variables: x-coordinate, y-coordinate, heading direction, pausing state, step length, and change in heading direction. Continuous variables were discretized using quantile-based symbolic encoding as implemented in the package (default settings) (69). Analyses were conducted with a history length of 1 time step (0.2 s). We evaluated longer lags (up to 10 steps) and found they did not qualitatively affect results. Net transfer entropy was defined as: T_anterior→posterior_ − T_posterior→anterior_ (30, 71), where positive values indicate predominant information flow from anterior to posterior. We tested whether net transfer entropy was greater than zero using one-sided one-sample t-tests.

### Model analysis

We formulated a discrete-time dynamical system that encodes order-dependent, role-asymmetric interactions between anterior and posterior components. Anterior trajectories function as generative components, while coordination is determined by a posterior response mechanism to the time integration of the anterior history. Importantly, the model architecture preserved anterior independence while maintaining cohesion, paralleling functional differentiation between anterior and posterior components, and permits systematic variation in role differentiation. In contact-based communication, body shape cannot be ignored. Thus, there is a difference between the anterior position **r**_**L**_(*t*) (e.g., the body center for the leader) and the actual point where the contact happens (e.g., the leader’s posterior tip of the body). We defined the point of interaction as an anterior proxy, denoted **r**_**P**_(*t*), to distinguish it from the anterior position, which governs locomotor control. A posterior component detects the proxy **r**_**P**_(*t*) rather than the body center, and evaluates the discrepancy between **r**_**P**_(*t*) and its own pre-update position **r**_**F**_(*t*).

The proxy is defined through a recursive temporal weighting of past leader positions, where λcontrols the contribution of the current leader position:

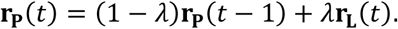

The follower first evaluates the error before updating its own motion

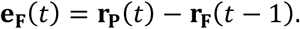

Finally, the follower’s position is updated according to this error. In this study, we consider the simplest realization of position updates, where the follower immediately moves to the proxy position:

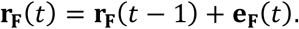

Thus, in this study, the follower position and proxy position became the same; these three equations can be simply described as

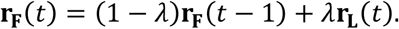

Further studies should consider a more natural implementation of locomotion. The proxy (and the follower) has exponentially weighted moving averages, so it possesses geometrically decaying memory governed by λ.

We tested the effect of λ, by generating the posterior trajectories with empirical anterior trajectories. For each anterior trajectory, we generated posterior trajectories across the range of λ from 0 to 1 in every 0.01. Then, we examined which λ value can create a similar empirical posterior trajectory. We obtained the root-mean-square errors (RMSEs) of the x- and y-coordinates across the trajectory while they were tandem running (within two body lengths, as in the above analysis). Then, we obtained the best fit λ for each trajectory. We compared the value of the best-fitted λ between tandem running pairs and single-termite coordination using t-tests for each species.

We used the global mean of the best-fitted λ (0.38) to obtain the fitted posterior trajectories to all empirical anterior trajectories. We investigated whether these model-generated trajectories exhibit patterns similar to those in empirical posterior trajectories. We used the same approach described above for empirical analysis. The major parts of the evolutionary simulation were constructed with C++, with the *Rcpp* v1.0.14 package integration (72).

## Acknowledgment

We would like to thank the Auburn University Social Insects Shared meeting for helpful comments. This study is supported by the USDA National Institute of Food and Agriculture, Hatch project number 7007938.

## Author contributions

N.M.: Conceptualization, Data Curation, Formal analysis, Methodology, Investigation, Validation, Visualization, Writing – original draft

K.S.: Conceptualization, Methodology, Investigation, Validation, Writing – review & editing

## Data accessibility

The data and codes for this study are available at https://github.com/nobuaki-mzmt/tandem_model. The accepted version will be deposited in Zenodo to obtain DOI.

## References

1. G. A. Cooper, S. A. West, Division of labour and the evolution of extreme specialization. Nature Ecology & Evolution 2, 1161–1167 (2018).

2. J. J. Boomsma, R. Gawne, Superorganismality and caste differentiation as points of no return: how the major evolutionary transitions were lost in translation. Biological Reviews 93, 28–54 (2018).

3. A. E. Emerson, Social coordination and the Superorganism. The American Midland Naturalist 21, 182–209 (1939).

4. B. R. Johnson, T. A. Linksvayer, Deconstructing the Superorganism: Social Physiology, Groundplans, and Sociogenomics. The Quarterly Review of Biology 85, 57–79 (2010).

5. C. R. Ward, F. Gobet, G. Kendall, Evolving Collective Behavior in an Artificial Ecology. Artificial Life 7, 191–209 (2001).

6. G. Theraulaz, E. Bonabeau, A Brief History of Stigmergy. Artificial Life 5, 97–116 (1999).

7. J. E. Herbert-Read, Understanding how animal groups achieve coordinated movement. Journal of Experimental Biology 219, 2971–2983 (2016).

8. N. T. Ouellette, A physics perspective on collective animal behavior. Physical Biology 19, 021004 (2022).

9. M. H. Dickinson, et al., How Animals Move: An Integrative View. Science 288, 100–106 (2000).

10. J. Chen, W. O. Friesen, T. Iwasaki, Mechanisms underlying rhythmic locomotion: body–fluid interaction in undulatory swimming. Journal of Experimental Biology 214, 561–574 (2011).

11. M. R. Greaney, E. S. Heckscher, M. T. Kaufman, Multiple scales of coordination along the body axis during Drosophila larval locomotion. J. Neurosci. (2026). 10.1523/JNEUROSCI.1623-25.2026.

12. A. Gomez-Marin, G. J. Stephens, M. Louis, Active sampling and decision making in Drosophila chemotaxis. Nature Communications 2, 441 (2011).

13. T. J. Prescott, M. E. Diamond, A. M. Wing, Active touch sensing. Philosophical Transactions of the Royal Society B: Biological Sciences 366 (2011).

14. A. G. Khan, M. Sarangi, U. S. Bhalla, Rats track odour trails accurately using a multi-layered strategy with near-optimal sampling. Nature Communications 3, 703 (2012).

15. K. C. Catania, Stereo and serial sniffing guide navigation to an odour source in a mammal. Nature Communications 4, 1441 (2013).

16. A. S. Machado, D. M. Darmohray, J. Fayad, H. G. Marques, M. R. Carey, A quantitative framework for whole-body coordination reveals specific deficits in freely walking ataxic mice. eLife 4, e07892 (2015).

17. I. D. Couzin, J. Krause, N. R. Franks, S. A. Levin, Effective leadership and decision-making in animal groups on the move. Nature 433, 513–516 (2005).

18. J. L. Harcourt, T. Z. Ang, G. Sweetman, R. A. Johnstone, A. Manica, Social feedback and the emergence of leaders and followers. Current Biology 19, 248–252 (2009).

19. J. W. Jolles, A. J. King, S. S. Killen, The Role of Individual Heterogeneity in Collective Animal Behaviour. Trends in Ecology & Evolution 35, 278–291 (2020).

20. A. A. Fernandez, J. L. Deneubourg, On following behaviour as a mechanism for collective movement. Journal of Theoretical Biology 284, 7–15 (2011).

21. X. G. Hou, D. J. Siveter, R. J. Aldridge, D. J. Siveter, Collective behavior in an early Cambrian arthropod. Science 322, 224 (2008).

22. J. T. Costa, T. D. Fitzgerald, A. Pescador-Rubio, J. Mays, D. H. Janzen, Social Behavior of Larvae of the Neotropical Processionary Weevil Phelypera distigma (Boheman) (Coleoptera: Curculionidae: Hyperinae). Ethology 110, 515–530 (2004).

23. P. Collard, Processionary Caterpillars at the Edge of Complexity. Artificial Life 30, 171–192 (2024).

24. W. Herrnkind, Queuing Behavior of Spiny Lobsters. Science 164, 1425–1427 (1969).

25. N. Mizumoto, C. R. Reid, Ant and termite collective behavior: Group-level similarity arising from individual-level diversity. Ecological Research 39, 646–658 (2024).

26. K. Tsuji, T. Ishikawa, Some observations of the caravaning behavior in the musk shrew (Suncus murinus). Behaviour 90, 167–183 (2008).

27. W. L. Nutting, “Flight and colony foundation.” in Biology of Termites, K. Krishna, F. M. Weesner, Eds. (Academic Press, 1969), pp. 233–282.

28. N. Mizumoto, S. Dobata, Adaptive switch to sexually dimorphic movements by partner-seeking termites. Science Advances 5, eaau6108 (2019).

29. N. Mizumoto, S. Reiter, Maintaining tandem movement cohesion through antennal movements in termites. Journal of The Royal Society Interface 22, 20250487 (2025).

30. G. Valentini, N. Mizumoto, S. C. Pratt, T. P. Pavlic, S. I. Walker, Revealing the structure of information flows discriminates similar animal social behaviors. eLife 9, e55395 (2020).

31. N. Mizumoto, T. Bourguignon, Light alters activity but does not disturb tandem coordination of termite mating pairs. Ecological Entomology 48, 145–153 (2022).

32. A. K. Raina, J. M. Bland, J. C. Dickens, Y. I. Park, B. Hollister, Premating behavior of dealates of the Formosan subterranean termite and evidence for the presence of a contact sex pheromone. Journal of Insect Behavior 16, 233–245 (2003).

33. N. Mizumoto, S. B. Lee, G. Valentini, T. Chouvenc, S. C. Pratt, Coordination of movement via complementary interactions of leaders and followers in termite mating pairs. Proceedings of the Royal Society B: Biological Sciences 288, 20210998 (2021).

34. T. Vicsek, A. Czirk, E. Ben-Jacob, I. Cohen, O. Shochet, Novel type of phase transition in a system of self-driven particles. Physical Review Letters 75, 1226–1229 (1995).

35. C. W. Reynolds Flocks, herds and schools: A distributed behavioral model. ACM SIGGRAPH Computer Graphics 21, 25–34 (1987).

36. I. D. Couzin, J. Krause, R. James, G. D. Ruxton, N. R. Franks, Collective memory and spatial sorting in animal groups. Journal of Theoretical Biology 218, 1–11 (2002).

37. D. Kong, K. Xue, P. Wang, Collective queuing motion of self-propelled particles with leadership and experience. Applied Mathematics and Computation 476, 128782 (2024).

38. T. E. Will, Flock Leadership: Understanding and influencing emergent collective behavior. The Leadership Quarterly 27, 261–279 (2016).

39. E. Cristiani, N. Loy, M. Menci, A. Tosin, Kinetic description and macroscopic limit of swarming dynamics with continuous leader–follower transitions. Mathematics and Computers in Simulation 228, 362–385 (2025).

40. M. Ballerini, et al., Interaction ruling animal collective behavior depends on topological rather than metric distance: evidence from a field study. Proceedings of the National Academy of Sciences of the United States of America 105, 1232–1237 (2008).

41. M. M. G. Sosna, et al., Individual and collective encoding of risk in animal groups. Proceedings of the National Academy of Sciences of the United States of America 116, 20556–20561 (2019).

42. J. E. Kowalko, et al., Loss of schooling behavior in cavefish through sight-dependent and sight-independent mechanisms. Current Biology 23, 1874–1883 (2013).

43. P. J. Gullan, P. S. Cranston, The Insects: An Outline of Entomology (John Wiley & Sons, 2014).

44. J. M. Rieser, et al., Geometric phase predicts locomotion performance in undulating living systems across scales. Proceedings of the National Academy of Sciences of the United States of America 121, e2320517121 (2024).

45. O. D. Broekmans, J. B. Rodgers, W. S. Ryu, G. J. Stephens, Resolving coiled shapes reveals new reorientation behaviors in C. elegans. eLife 5, e17227 (2016).

46. J. Gray, Directional control of fish movement. Proc Biol Sci 113, 115–125 (1933).

47. T. D. Fitzgerald, Role of trail pheromone in foraging and processionary behavior of pine processionary caterpillars Thaumetopoea pityocampa. Journal of Chemical Ecology 29, 513–532 (2003).

48. M. Nagy, Z. Ákos, D. Biro, T. Vicsek, Hierarchical group dynamics in pigeon flocks. Nature 464, 890–893 (2010).

49. J. Krause, D. Hoare, S. Krause, C. K. Hemelrijk, D. I. Rubenstein, Leadership in fish shoals. Fish and Fisheries 1, 82–89 (2000).

50. B. Pettit, Z. Ákos, T. Vicsek, D. Biro, Speed Determines Leadership and Leadership Determines Learning during Pigeon Flocking. Current Biology 25, 3132–3137 (2015).

51. L. J. N. Brent, et al., Ecological knowledge, leadership, and the evolution of menopause in killer whales. Current Biology 25, 746–750 (2015).

52. N. R. Franks, T. O. Richardson, Teaching in tandem-running ants. Nature 439, 153 (2006).

53. T. Sasaki, L. Danczak, B. Thompson, T. Morshed, S. C. Pratt, Route learning during tandem running in the rock ant Temnothorax albipennis. The Journal of Experimental Biology 223, jeb221408 (2020).

54. T. O. Richardson, P. A. Sleeman, J. M. McNamara, A. I. Houston, N. R. Franks, Teaching with evaluation in ants. Current Biology 17, 1520–1526 (2007).

55. N. Mizumoto, et al., Functional and mechanistic diversity in ant tandem communication. iScience 26, 106418 (2023).

56. L. Conradt, T. J. Roper, Consensus decision making in animals. Trends in Ecology and Evolution 20, 449–456 (2005).

57. K. Tunstrøm, et al., Collective states, multistability and transitional behavior in schooling fish. PLOS Computational Biology 9, e1002915 (2013).

58. C. Heins, et al., Collective behavior from surprise minimization. Proceedings of the National Academy of Sciences of the United States of America 121, e2320239121 (2024).

59. C. Huang, F. Ling, E. Kanso, Collective phase transitions in confined fish schools. Proceedings of the National Academy of Sciences of the United States of America 121, e2406293121 (2024).

60. M. Guttenberg, S. Sripad, V. Viswanathan, Evaluating the Potential of Platooning in Lowering the Required Performance Metrics of Li-Ion Batteries to Enable Practical Electric Semi-Trucks. ACS Energy Letters 2, 2642–2646 (2017).

61. S. Feng, et al., String stability for vehicular platoon control: Definitions and analysis methods. Annual Reviews in Control 47, 81–97 (2019).

62. O. Yamanaka, R. Takeuchi, UMATracker: An intuitive image-based tracking platform. Journal of Experimental Biology 221, 1–24 (2018).

63. T. D. Pereira, et al., SLEAP: A deep learning system for multi-animal pose tracking. Nature Methods 19, 486–495 (2022).

64. R Core Team, R: A language and environment for statistical computing. Ver 4.5.2. (2025). Deposited 2025.

65. N. Mizumoto, S.-B. Lee, T. Chouvenc, The strength of sexual signals predicts same-sex pairing in two Coptotermes termites. Behavioral Ecology 35, arae067 (2024).

66. D. M. Bates, M. Maechler, Package “lme4” Linear Mixed-Effects Models using “Eigen” and S4. Journal of Statistical Software 67, 1–48 (2015).

67. F. Bartumeus, et al., Foraging success under uncertainty: search tradeoffs and optimal space use. Ecology Letters 19, 1299–1313 (2016).

68. T. Hothorn, Multcomp: Simultaneous inference in general parametric models. R package version 1, 4 (2020).

69. S. Behrendt, T. Dimpfl, F. J. Peter, D. J. Zimmermann, RTransferEntropy — Quantifying information flow between different time series using effective transfer entropy. SoftwareX 10, 100265 (2019).

70. T. Schreiber, Measuring information transfer. Physical Review Letters 85, 461–464 (2000).

71. M. Porfiri, Inferring causal relationships in zebrafish-robot interactions through transfer entropy: a small lure to catch a big fish. Animal Behavior and Cognition 5, 341–367 (2018).

72. D. Eddelbuettel, J. J. Balamuta, Extending R with C++: A Brief Introduction to Rcpp. The American Statistician 72, 28–36 (2018).

