## Supplementary material for "Body-like coordination emerges in paired termite locomotion": Table S1-2 Figure S1-12 Legends Video S1

ORCID: NM: 0000-0002-6731-8684; SK: 0000-0002-3855-8048

**Table of contents**

**Table S1-2**
**Figure S1-12**
**Legends Video S1**

**Table S1. Source of all data used in the study**

| Species | Behavior | Source | Location | Year | N | Arena (mm) | Duration (min) | Tracking | Body length |
|---|---|---|---|---|---|---|---|---|---|
| *C. formosanus* | Tandem | (1) | Wakayama, JP | 2017 | 17 | 145 | 60 | UMATracker | (2) |
| | Tandem | (3) | Florida, USA | 2020 | 10 | 140 | 30 | UMATracker | (4) |
| | Tandem | (5) | Okinawa, JP | 2022 | 10 | 90 | 30 | SLEAP | (5) |
| | Solo | (5) | Okinawa, JP | 2022 | 20 | 90 | 15 | SLEAP | (5) |
| *R. speratus* | Tandem | (1) | Kyoto, JP | 2017 | 20 | 145 | 60 | UMATracker | (2) |
| | Tandem | (5) | Fukui/Ishikawa, JP | 2022 | 20 | 45 | 15 | SLEAP | (5) |
| | Solo | (5) | Okinawa, JP | 2022 | 40 | 45 | 15 | SLEAP | (5) |
| *T. rugatulus* | Tandem | (2) | Arizona, USA | 2018 | 20 | NA | 10 | UMATracker | (2) |
| *D. indicum* | Tandem | (6) | Okinawa, JP | 2020 | 15 | NA | 10 | UMATracker | (6) |


1. N. Mizumoto, S. Dobata, Adaptive switch to sexually dimorphic movements by partner-seeking termites. *Science Advances* **5**, eaau6108 (2019).
2. G. Valentini, N. Mizumoto, S. C. Pratt, T. P. Pavlic, S. I. Walker, Revealing the structure of information flows discriminates similar animal social behaviors. *eLife* **9**, e55395 (2020).
3. N. Mizumoto, S. B. Lee, G. Valentini, T. Chouvenc, S. C. Pratt, Coordination of movement via complementary interactions of leaders and followers in termite mating pairs. *Proceedings of the Royal Society B: Biological Sciences* **288**, 20210998 (2021).
4. N. Mizumoto, S.-B. Lee, T. Chouvenc, The strength of sexual signals predicts same-sex pairing in two *Coptotermes* termites. *Behavioral Ecology* **35**, arae067 (2024).
5. N. Mizumoto, S. Reiter, Maintaining tandem movement cohesion through antennal movements in termites. *Journal of The Royal Society Interface* **22**, 20250487 (2025).
6. N. Mizumoto, *et al.*, Functional and mechanistic diversity in ant tandem communication. *iScience* **26**, 106418 (2023).

**Table S2.** Best fitted λ for each species and data type.

| Species | Type | n | λ, Mean (95% CI) |
|---|---|---|---|
| *C. formosanus* | Solo | 20 | 0.393 [0.364–0.423] |
| | Tandem | 37 | 0.369 [0.356–0.382] |
| *R. speratus* | Solo | 40 | 0.392 [0.363–0.421] |
| | Tandem | 40 | 0.368 [0.349–0.387] |
| Global | All | 137 | 0.379 [0.367–0.390] |

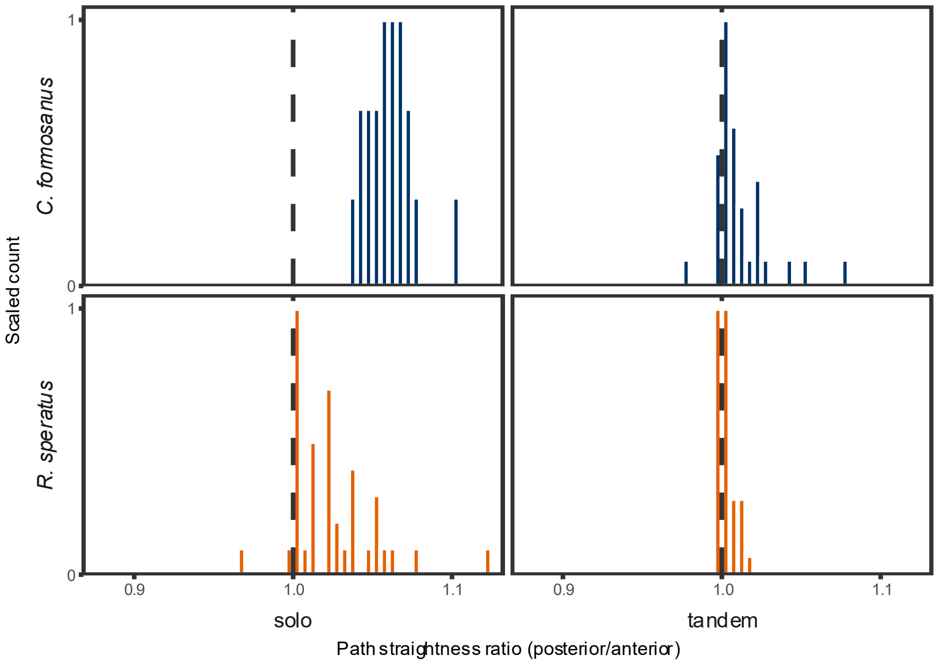


**Figure S1.** Path straightness ratio comparison between anterior and posterior components. One-sample t-test showed that all of these are greater than 1 ($P < 0.002$), where the normality was not violated in the case of the solo of *C. formosanus* (Shapiro test $P = 0.283$). The nonparametric Wilcoxon signed-rank test yielded the same conclusion ($P < 0.01$ for all).

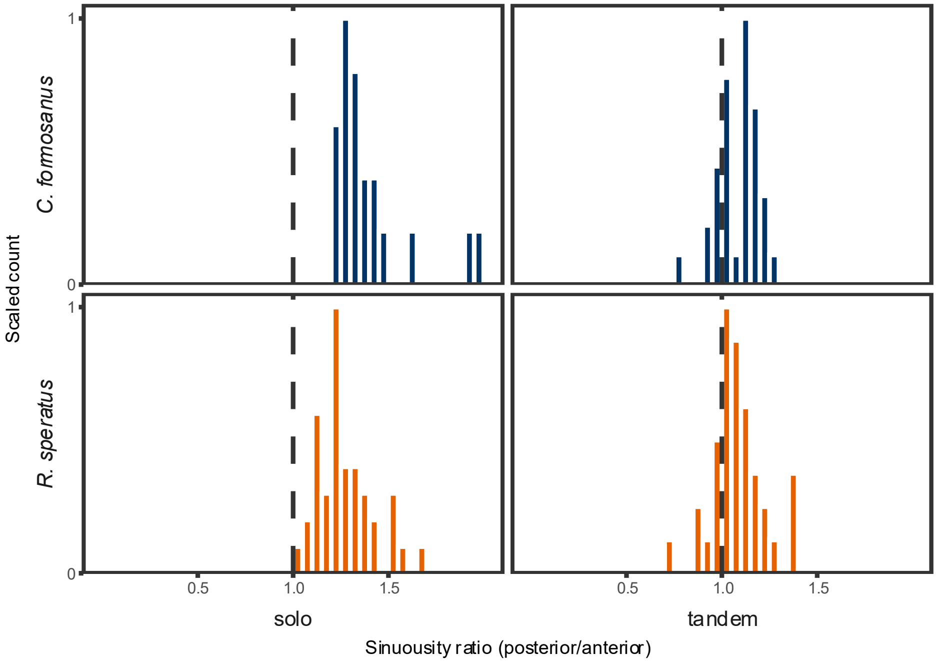


**Figure S2.** Sinuousity ratio comparison between anterior and posterior components. Sinuousity was measured as the mean turning angles of each successive frame. One-sample t-test showed that all of these are greater than 1 ($P < 0.002$), where the normality was not violated in the case of the tandem of *C. formosanus* (Shapiro test $P = 0.950$). The nonparametric Wilcoxon signed-rank test yielded the same conclusion ($P < 0.002$ for all).

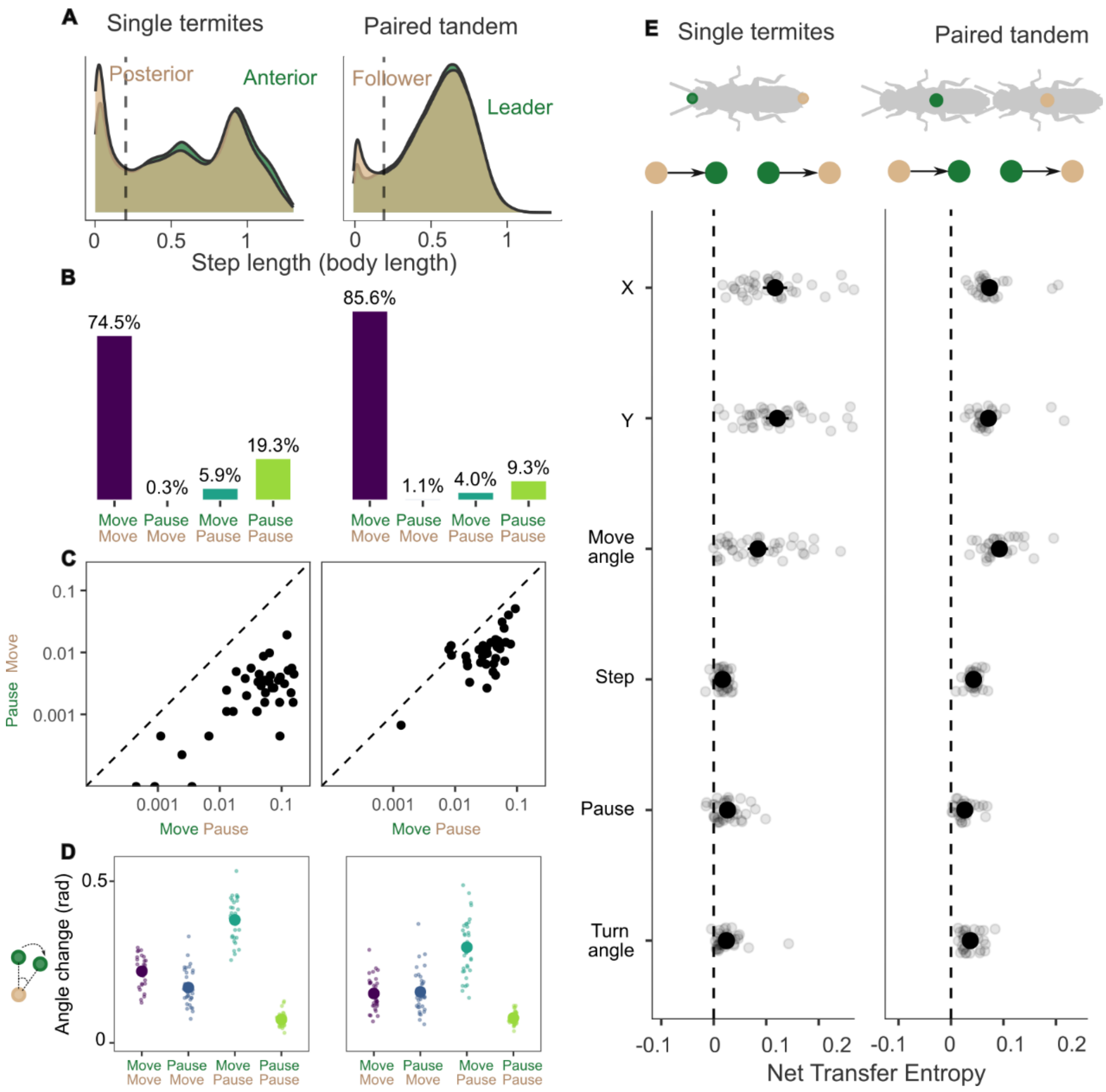


**Figure S3**. Movement coordination in paired and single termites shares the same division of labor and directional control. Data are from *R. speratus*. (A) Step length distributions show shorter and more intermittent movement in posterior parts (or followers) compared to anterior parts (or leaders), consistent with stabilizing versus exploratory roles (see Fig. S5 for breakdown). Dashed lines indicate the pausing threshold (0.2 body length). (B) Frequencies of movement states show strong asymmetry: posterior/follower pausing while anterior/leader moves occur more frequently than the reverse. (C) This asymmetry is consistent across individuals and pairs. (D) Turning depends on movement state and is greatest when the anterior/leader moves while the posterior/follower pauses, indicating exploration during posterior waiting. Different letters indicate significant differences (LMM; Tukey HSD, $P < 0.001$). (E) Net transfer entropy (TE_ante→post − TE_post→ante) shows consistent directional information flow from anterior to posterior or leader to follower across kinematic and behavioral variables. * indicates the statistical significance that net transfer entropy is larger than 0 (one-sample t-test).

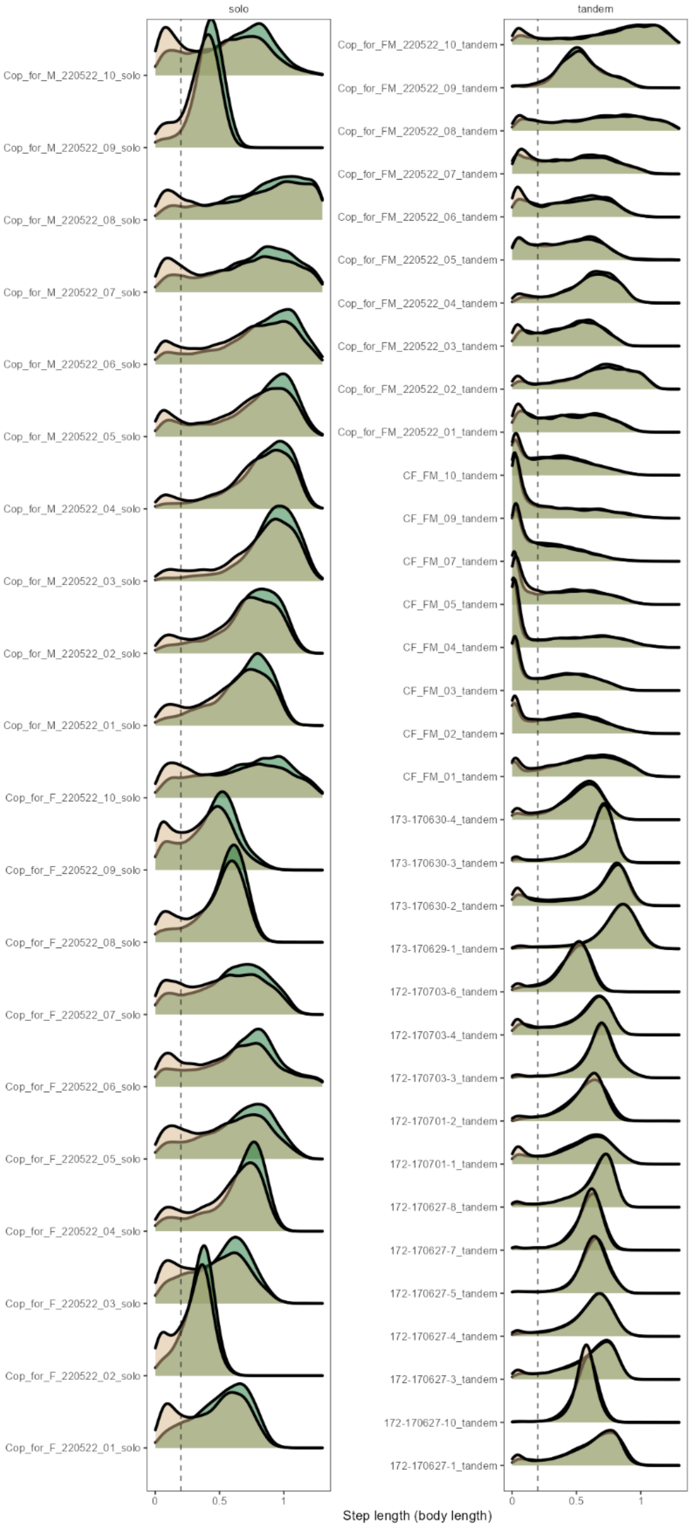


**Figure S4.** Step length distribution for each individual or pair in *C. formosanus*. The left is for solo running, and the right is for tandem running.

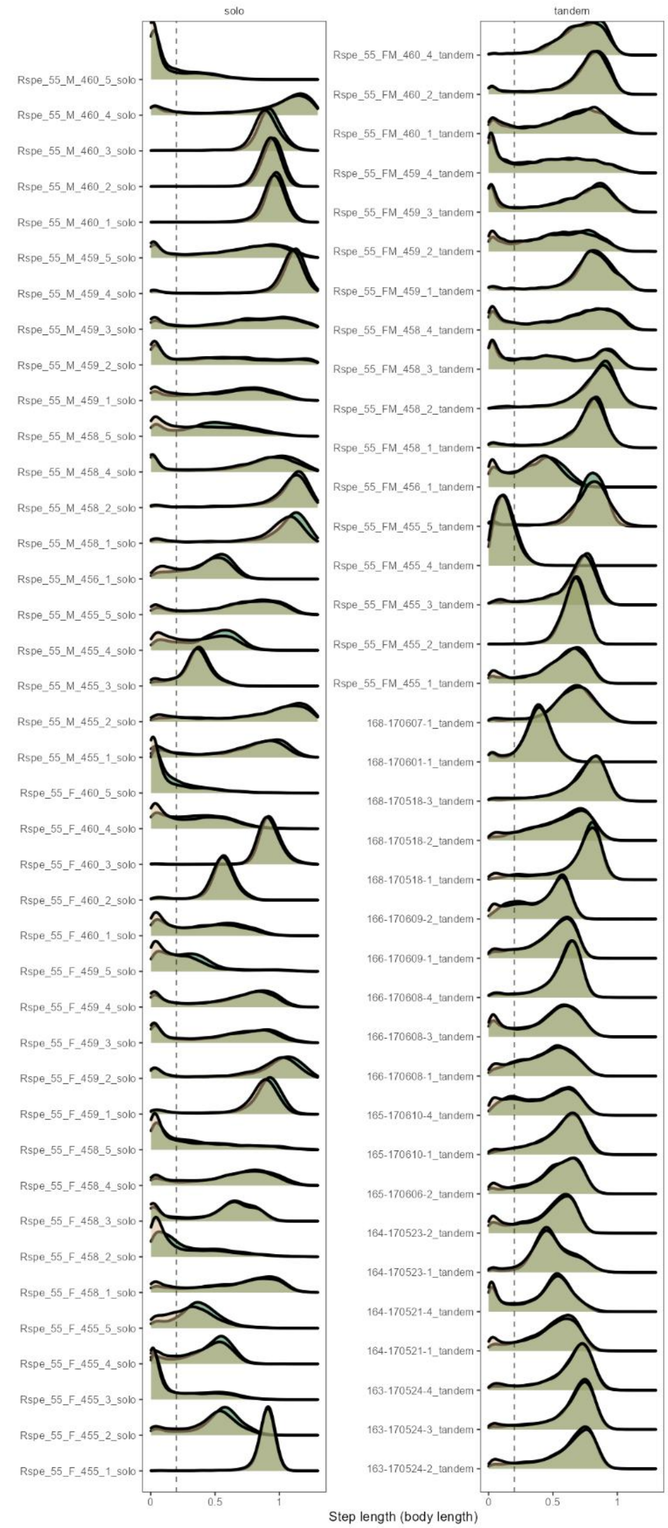


**Figure S5.** Step length distribution for each individual or pair in *R. speratus*. The left is for solo running, and the right is for tandem running.

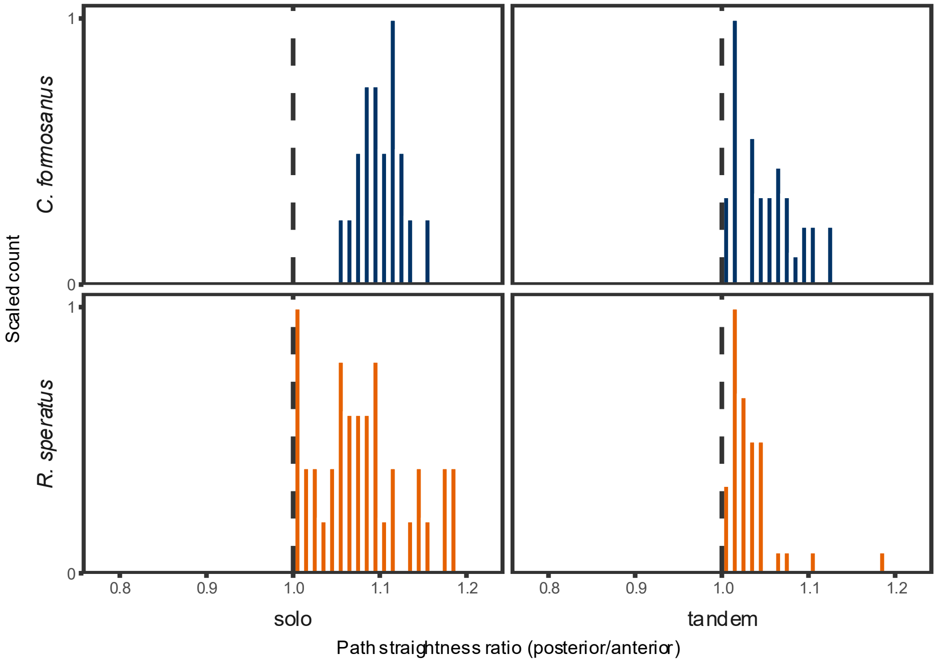


**Figure S6.** Path straightness ratio comparison between anterior and posterior components in model-fitted data. One-sample t-test showed that all of these are greater than 1 ($P < 0.001$), where the normality was not violated in the case of the solo datasets (Shapiro test $P > 0.05$). The nonparametric Wilcoxon signed-rank test yielded the same conclusion ($P < 0.001$ for all).

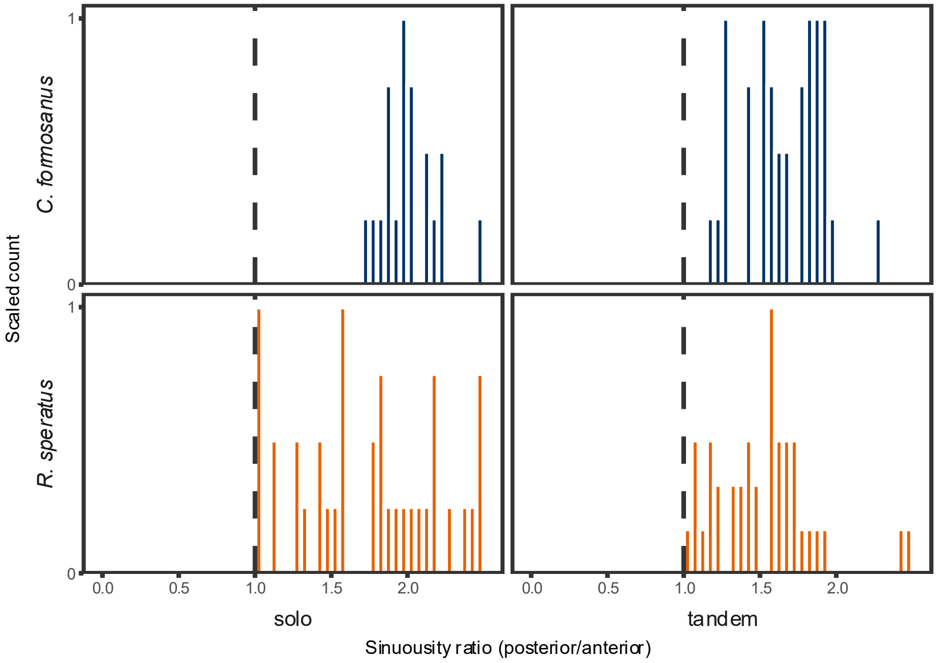


**Figure S7.** Sinuousity ratio comparison between anterior and posterior components in model-fitted data. Sinuousity was measured as the mean turning angles of each successive frame. The one-sample t-test showed that all of these were greater than 1 (P < 0.001), and normality was violated in the case of the tandem of *R. speratus* (Shapiro test, $P = 0.008$). The nonparametric Wilcoxon signed-rank test yielded the same conclusion ($P < 0.001$ for all).

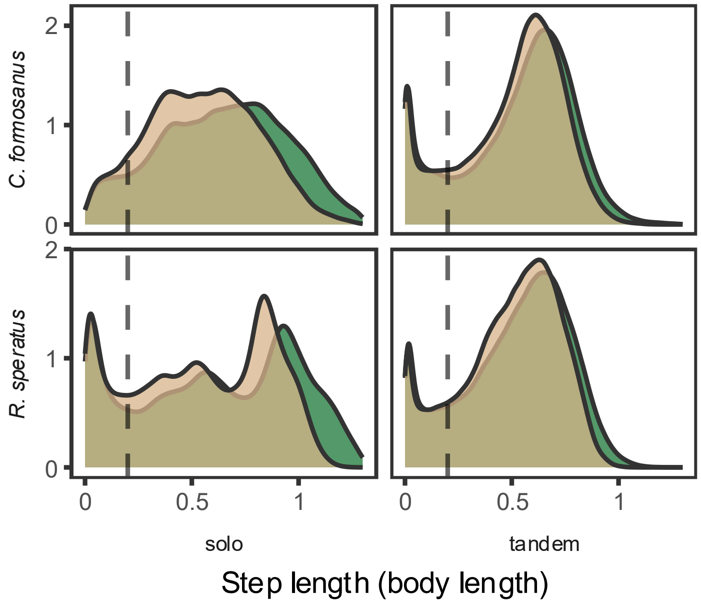


**Figure S8.** Step length distribution for model-fitted data. Green is the empirical anterior trajectory, while orange is the model-fitted posterior trajectory (λ = 0.38).

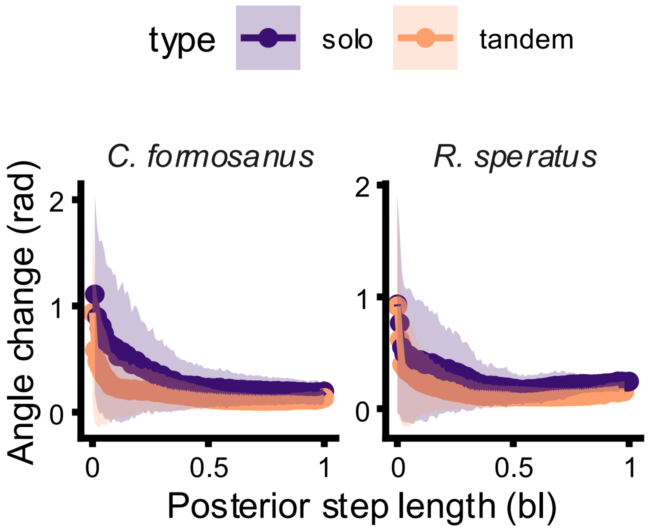


**Figure S9.** The relationship between posterior step length and changes in the angle between posterior and anterior positions (λ = 0.38).

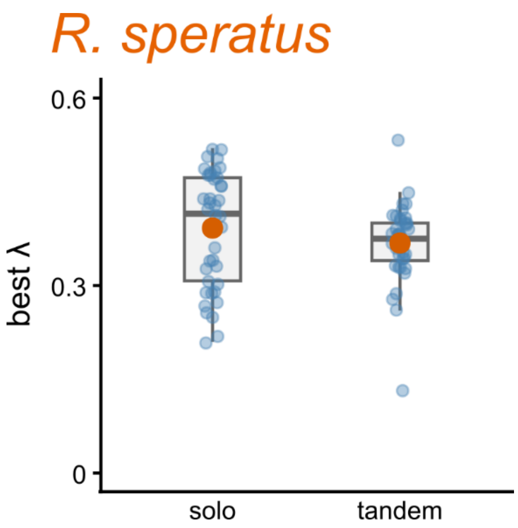


**Figure S10.** Comparison of the best fitted λ for each trajectory of single termites and paired termites in *R. speratus*. There was no significant difference between within-individual and across-individual coordination (t-test, $t_{68.2} = 1.37$, $P = 0.18$).

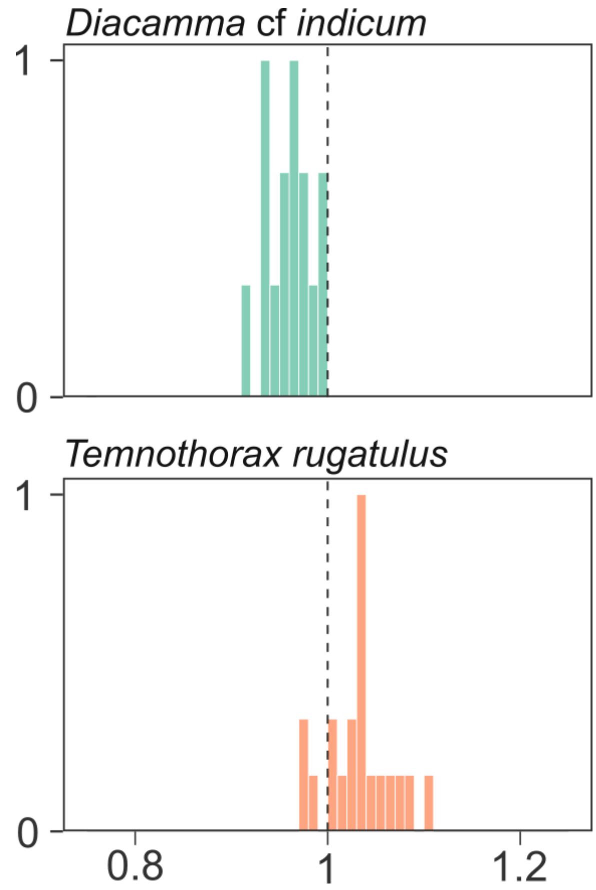


**Figure S11.** Comparison of the leader and follower trajectory length in ants. In both species, the mean values were biased from 1, with *D. indicum* significantly smaller than 1 (t-test, $t_{14} = -6.76$, $P < 0.001$) and *T. rugatulus* significantly larger than 1 (t-test, $t_{19} = 4.13$, $P < 0.001$). Neither violated the assumption of normality (Shapiro-Wilk Test, $P > 0.69$).

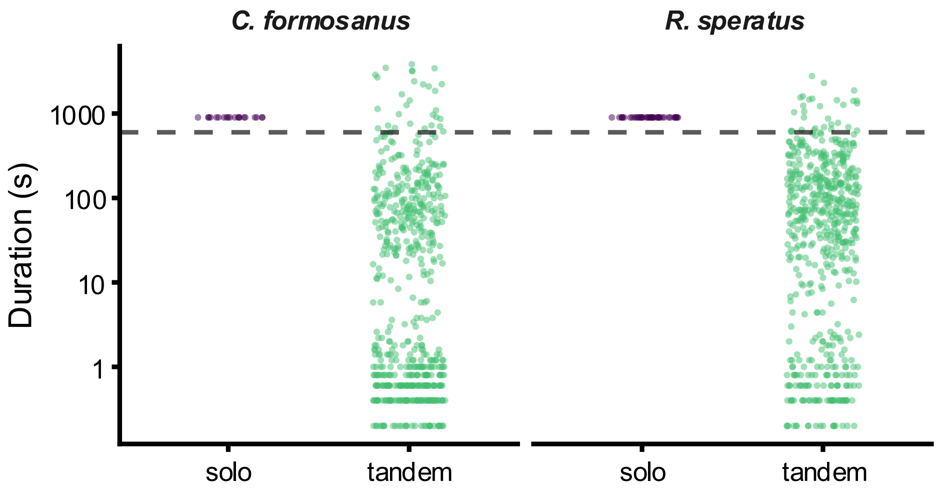


**Figure S12.** Distribution of the duration of tandem running (defined as within two body lengths). The dashed line is the threshold for inclusion in the transfer entropy analysis.

**Legends for Supplementary Videos**

**Video S1.** Comparison of movement trajectories between paired termites (leader and follower) and single termites (head tip and abdomen tip). The color indicates the anterior or posterior components.
